# Highly responsive single-fluorophore indicator to explore lactate dynamics in high calcium environments

**DOI:** 10.1101/2020.10.01.322404

**Authors:** A. Galaz, PY. Sandoval, I. Soto-Ojeda, H. Hertenstein, J. Schweizer, S. Schirmeier, L.F Barros, A. San Martín

## Abstract

Lactate is an energy substrate and intercellular signaling molecule with multiple bodily functions. Lactate has physiological roles in neurogenesis, axon integrity, memory consolidation, immune response, exercise, adipose tissue lipolysis, etc, and is involved in inflammation, cancer and neurodegeneration. The FRET lactate indicator Laconic has been instrumental in the discovery of mechanisms involved in neurometabolic coupling, and has advanced the understanding of lactate transport, glycolysis and mitochondrial physiology. However, the low fluorescent response and the complex saturation kinetics of Laconic limit its use for high-throughput screening and quantitation. Using the bacterial periplasmic binding protein TTHA0766 from *Thermus thermophilus*, we have now developed the first single-fluorophore indicator for lactate. The sensor exhibited an intensiometric fluorescence increase of ΔF/F_0_ 3.0 and a single binding site with a K_D_ of 293 μM. The fluorescence is not affected by other monocarboxylates or pH. However, it is sensitive to Ca^2+^ in the nanomolar range. Targeting of the sensor to the endoplasmic reticulum revealed that this organelle presents a high permeability for lactate. The functionality of the sensor in living tissue is demonstrated in the brain of *Drosophila melanogaster* larvae. This indicator, which we have termed CanlonicSF, is well suited to explore lactate dynamics in environments with micromolar Ca^2+^ or higher, such as the endoplasmic reticulum and the extracellular space.

## INTRODUCTION

Lactate is a monocarboxylate that was long considered a waste product of hypoxic metabolism. However, recent evidence indicates that under physiological conditions lactate is an energy substrate and a signaling molecule. Lactate plays a role in long-term memory consolidation, maintaining <u>L</u>ong-<u>T</u>erm <u>P</u>otentiation (LTP)^1, 2^, and neurogenesis^3^. It also exerts neuroprotective effects and is necessary for axon integrity^4–8^. Furthermore, lactate plays a pivotal role in the pathology of inflammation, cancer, neurodegeneration, and microbial infection.

Lactate detection has been performed by enzymatic reaction followed by photometric or amperometric procedures and spectrometry techniques such as <u>N</u>uclear <u>M</u>agnetic <u>R</u>esonance (MNR) or <u>H</u>igh-<u>P</u>ressure <u>L</u>iquid <u>C</u>hromatography coupled with <u>M</u>ass <u>S</u>pectrometry (HPLC-MS). To monitor lactate dynamics with improved spatial/temporal resolution in living systems, we developed a <u>F</u>örster <u>R</u>esonance <u>E</u>nergy <u>T</u>ransfer (FRET)-based indicator called Laconic^9^. This tool has been instrumental in the discovery of mechanisms involved in neurometabolic coupling^10–14^, identification of new <u>m</u>ono<u>c</u>arboxylate <u>t</u>ransporters (MCT)^15^ and channels^16^, improved monocarboxylate transporter characterization^17^, development of high-throughput methodologies to detect mitochondrial toxicity^18^, and a complete set of methodologies to evaluate lactate dynamics in intact systems^19^.

FRET indicators are excellent research tools since they are ratiometric and pH resistant. However, these indicators have a low fluorescent response and are difficult to target to organelles such as mitochondria, therefore hampering lactate monitoring with sub-cellular resolution. Laconic ‒ the only genetically encoded indicator for lactate currently available – binds to lactate with two apparent binding sites, which determines a wide dynamic range spanning from 1 μM to 10 mM^9^. The tradeoff is that accurate quantification is difficult, especially *in vivo* where the experimental control is hard to achieve. A highly responsive lactate indicator ‒ with much higher fluorescence response and one site lactate dose-response curve ‒ could, thus, be valuable for the development of more sensitive high-throughput methodologies and accurate lactate quantification.

Highly responsive genetically encoded indicators can be engineered inserting a <u>c</u>ircularly <u>p</u>ermuted <u>G</u>reen <u>F</u>luorescent <u>P</u>rotein (cpGFP) into the backbone of the ligand-binding moiety^20, 21^. The interface between the fluorescent reporter and the binding ligand module is close to the chromophore because the natural terminus of the GFP molecule is fused to a peptide linker, whereas a new terminus will be formed in a region close to the chromophore, exposing it to the solvent^22^. As a result, ligand-induced structural rearrangements are transduced to the chromophore surroundings, modulating its pK_a_, solvent access, absorption and quantum yield, and therefore chromophore brigthness^21, 23, 24^. Typically, the first generation of this so-called single-fluorophore indicator displays between a 100 % and 300 % change in fluroescence^20^. This is higher than FRET indicators, which show a lower change in fluorescence, ranging from 10 to 80 %^9, 25–27^. Single-fluorophore indicators also present advantages in comparison with intramolecular FRET-based indicators: they are easier to target to sub-cellular localizations, linkers can be easily engineered to render optimized versions, permits multiplexing measurements, and allowing signal detection with a user-friendly and widely available optical setup.

In this article, we introduce a highly responsive GFP-based single-fluorophore calcium-dependent lactate indicator, CanlonicSF (<u>Ca</u>lcium-a<u>n</u>d-<u>L</u>actate <u>Op</u>tical <u>N</u>ano-<u>I</u>ndicator from <u>C</u>ECs <u>S</u>ingle <u>F</u>luorophore). This indicator is based on the <u>P</u>eriplasmic <u>B</u>inding <u>P</u>rotein (PBP) TTHA0766 from *Thermus thermophilus* strain HB8, for which the calcium-lactate bound crystallographic structure is known^28^. We demonstrate that a single insertion point of a cpGFP is permissive and produces a functional and specific lactate indicator at micromolar calcium concentration. CanlonicSF permits the monitoring of lactate dynamics in mammalian systems with high spatial/temporal resolution and is suited for high calcium environments, such as within the endoplasmic reticulum (ER) and the extracellular space.

## MATERIAL AND METHODS

Standard reagents and inhibitors were acquired from Sigma or Merck.

### Construction of a lactate single-fluorophore indicator

*The TTHA0766* gene from *Thermus thermophilus* HB8 was synthesized without a periplasmic target sequence by GenScript and the plasmid encoding cpGFP was kindly provided by Wolf Frommer. Guided by the crystallographic structure of TTHA0766 sixteen sites were selected for cpGFP insertion ‒ flanked by two aminoacid linkers LE/TR – into the backbone of calcium-lactate binding PBP. These insertion sites were in external loops to avoid the hydrophobic protein core, which is critical for proper protein folding. Each DNA construct was made up of three parts: TTHA0766 was split into two fragments at the point where the cpGFP was inserted. Each of these three amplicons were amplified by Polymerase Chain Reaction (PCR) using the high-fidelity DNA polymerase KOD (Merck, Darmstadt, Germany). The primers used to amplify the three amplicons of the sixteen prototypes were designed with overlapping regions to perform overlapping PCR with primers which included attB1 and attB2 Gateway recombination sites at the forward and reverse primer, respectively. The resulting amplicon from overlapping PCR was cloned to entry plasmid pDONR 221 (ThermoFisher, Waltham, Massachusetts, USA), using BP recombination reaction using Gateway technology (ThermoFisher, Waltham, Massachusetts, USA). Each entry plasmid was recombined with a destination plasmid performing an LR recombination reaction, with a modified version of pcDNA3.1(−) (ThermoFisher, Waltham, Massachusetts, USA), which contained attR1 and attR2 recombination sites. The DNA sequence of all the resulting expression vectors were verified through DNA sequencing (AustralOMICs, UACh, Valdivia, Chile)

### Protein Purification

Protein extracts were taken from a cell line stably expressing CanlonicSF (HEK-024), which was generated by lentivirus transduction. Briefly, three 100 millimeters plates with clonal cells HEK024 were cultured at 37°C in 95% air/5% CO_2_ for 72 hours or until they reached 100 % confluency. Then the media culture was removed, and cells were washed twice with 10 ml of cold 1X PBS containing in mM, 176 NaCl, 2.6 KCl, 10 Na_2_HPO_4_, and 27.6 KH_2_PO_4_, pH 7.0. Next, a third wash with 1X PBS was used to detach the plated cells. Cells were centrifuged at 2500 rpm for 10 minutes. The cell pellet was resuspended in 500 μl of 1X PBS buffer and lysed by six freeze/thaw cycles. The lysate was diluted in 10 ml of 1X PBS buffer and filtered using an Amicon 50 kDa filter column (Merck, Darmstadt, Germany) to concentrate the extract to a final volume of 500 μL.

### Fluorescence Measurements

Purified proteins were resuspended in an intracellular buffer containing in mM: 10 NaCl, 130 KCl, 1.25 MgSO_4_, and 10 HEPES, pH 7.0. The purified extracts of CanlonicSF were excited at 485 ± 20 nm and the intensity of fluorescence emission of cpGFP was recorded at 530 ± 15 nm using the multiwell plate analyzer Synergy 2 (Biotek, Winooski, Vermont, USA). Excitation and emission spectra were obtained by collecting the emissions at 515 ± 15 nm using a multiwell plate reader EnVision (PerkinElmer, Waltham, Massachusetts, USA).

Imaging experiments in HEK293 and COS7 cells from the American Type Culture Collection (ATCC) were performed at 37°C in 95% air/5% CO2 in DMEM/F12 10% fetal bovine serum (HEK293) or DMEM high glucose with 10% fetal bovine serum (COS7). Cell lines were transfected at 60% confluency using Lipofectamine 3000 (Gibco), with an efficiency of > 60% and imaged at room temperature (22 - 25°C). Cells were superfused with a 95% air/5% CO2-gassed solution of the following composition (in mM): 112 NaCl, 1.25 CaCl2, 1.25 MgSO_4_, 1-2 glucose, 10 HEPES, 24 NaHCO_3_, 1 EGTA, 5 KCl, pH 7.4, using an upright Olympus FV1000 confocal microscope equipped with a 20x water immersion objective (N.A. 1.0) and a 440 nm solid-state laser. Times series images were taken every 10 seconds with 20x (NA 1.0) in XYT scan mode (scan speed: 156 Hz; 800 × 800-pixel; pinhole 800 μm).

Live imaging of *ex vivo* Drosophila brains was performed at room temperature with an inverted Leica Dmi8 SP8 confocal microscope (Leica Microsystems GmbH, Heidelberg). Times series images were taken every 5 seconds with a 20x HC PL APO objective (NA 0.75 CS2) in single plane mode (scan speed: 600 Hz; 1024 × 1024 pixel, 8 × 8 binning, 0.2 μm × 0.2 μM, pinhole 2 AU). Third instar larval brains were dissected in HL3−/− buffer (70 mM NaCl, 5 mM KCl, 20 mM MgCl_2_, 10 mM NaHCO_3_, 115 mM sucrose, 5 mM HEPES; pH 7.2; ca. 350 mOsm), directly attached to a Poly-D-Lysine coated glass coverslip (Sigma-Aldrich, P8920) and transferred onto a custom made flow-through chamber. Buffer flow was adjusted by a variable speed pump (neoLab Migge GmBH, Heidelberg, Germany) to achieve a constant current (4,5 ml/min). Buffer was aspirated by a custom-made vacuum pump using a commercial aquarium pump (Tetra Aquarium Air Pump, APS 50). Samples were excited at 440 nm, beam splitter DD488/552; emission was recorded at 495 - 547 nm. For ratiometric measurements the fluorescence intensity of a red fluorophore with matching expression pattern to the sensor was used. tdTomato was excited at: 552 nm and emission was detected between 566 – 637 nm.

### Transgenic *Drosophila melanogaster* generation

The flies harboring the sensor CanlonicSF were created by the following strategy: The coding sequence of the sensor has been amplified from pcDNA3.1 using the primers caccATGTTTTCTCCTCTGGCGG and CTAAAGGGAAAGCCCCTTG and subsequently inserted into a pENTR-vector using TOPO® cloning (ThermoFisher, Waltham, Massachusetts, USA), and afterwards into a pUASTattBrfa^29^ vector via LR recombination using Gateway technology (ThermoFisher, Waltham, Massachusetts, USA). The pUASTattBrfa vector allows Φ integrase-mediated recombination of the construct into the fly genome at landing site attp40. The resulting transgene was recombined to a UAS-tdtomato transgene^30^ to generate double transgenic flies.

### Data and statistical analysis

For data analysis, the fluorescent signal from a region of interest (ROI) from each cell was collected. Background subtraction was performed separately for each channel. Regression and statistical analyses were carried out using SigmaPlot (Jandel).

## RESULTS

### Development of the single-fluorophore lactate indicator

Building a single-fluorophore indicator requires the fusion of two modules, one to detect a ligand of interest and another to report the binding through fluorescence. Typically, the ligand-binding moieties that have been used for single-fluorophore indicators are transcription factors (TF) and periplasmic binding proteins (PBP) from bacteria^20^. To the best of our knowledge, there are two bacterial proteins that detect lactate with the affinity and specificity needed to develop genetically encoded indicators; LldR^31, 32^, and TTHA0766^28^. The first obvious candidate was the bacterial TF LldR, which had been successfully used to develop the first FRET-based indicator for lactate, Laconic^9^. Single-fluorophore prototypes using LldR failed, due to the severe pH sensitivity presented in all variants tested. Therefore, our next candidate was the crystalized PBP from *Thermus thermophilus* HB8, TTHA0766. The crystallographic structure of this PBP showed a coordinated lactate molecule, even though lactate was not added during the crystallization process ^28^. This suggested the presence of a high-affinity lactate binding site. Also, in the binding pocket, a calcium ion that was in direct contact with lactate was identified^28^, suggesting an important role for calcium in the binding and high affinity of TTHA0766 for this monocarboxylate. High-affinity ligand binding moieties are good candidates to develop single-fluorophore indicators. Insertion of fluorescence modules such as cpGFP into the backbone of the binding protein can produce a structural perturbation, decreasing its affinity for a given ligand. Physiological lactate spans the millimolar range, so a cpGFP insertion-induced affinity modification from the nanomolar to millimolar range is highly desirable.

Taking advantage of the available crystallographic structure of TTHA0766 from *Thermus thermophilus* HB8, we selected 16 sites in the backbone of TTHA0766 to test their suitability for cpGFP insertion (**Figure 1A**). These sites were selected based on their localization in external protein loops, to avoid the perturbation of the hydrophobic protein core and hydrophobic collapse during protein folding. To screen suitable lactate indicators, all 16 prototypes were expressed in HEK293 and exposed to pulses of lactate and pyruvate to evaluate their response. One prototype with the insertion point in the amino acid leucine in position 177 was permissive to cpGFP insertion (**Figure 1A and B**). This prototype presented a clear response to lactate and did not respond to pyruvate (**Figure 1B**). The rest of the prototypes showed a weak response to lactate and pyruvate.

**Figure 1.**
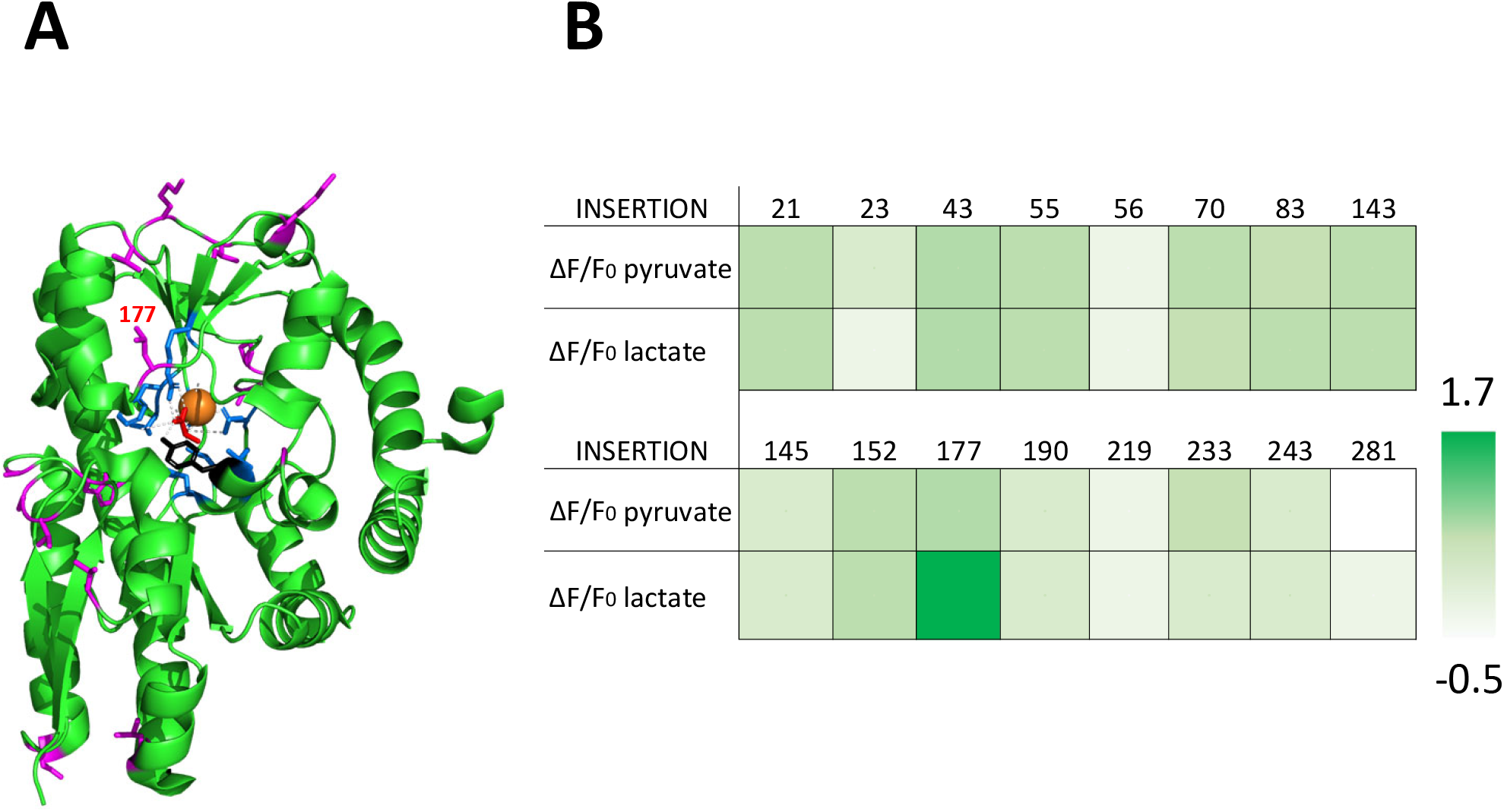
Design and Screening of a Single-Fluorophore Indicator for Lactate Based on the Periplasmic Protein from *Thermus thermophilus*, TTHA0766. A) Schematic representation of the crystallography structure of TTHA0766^28^ using PyMol. Lactate molecule in red, calcium cation in orange, amino acids of the binding pocket site are in blue, tyrosine residue mutated to generated CanlonicSF in black, and amino acids, where cpGFP was inserted, are highlighted in magenta. B) Heatmap of normalized fluorescence response to 5 mM lactate and pyruvate of a series of prototypes with different cpGFP insertion points expressed in HEK293 cells.

Typical PBPs present nanomolar affinity for their ligands^33^, below the physiological range in mammalian fluids. The insertion of a fluorescence module produces a perturbation in the chimeric protein that induces a decrease in affinity by two orders of magnitude, reaching the micromolar range. To decrease the affinity of this prototype further into the physiological range we performed alanine scanning of the aminoacid residues in close contact with lactate. The mutation of tyrosine 101 located in the lactate pocket binding site^28^ (**Figure 1A**) by alanine produces a low-affinity variant that we christened as CanlonicSF.

### *In vitro* characterization of CanlonicSF

Further, to characterize the behavior of CanlonicSF, we generated a HEK293 stably expressing the sensor and obtained protein extracts. First, a dose-response curve was performed to determine its dynamic range and dissociation constant. Exposure to increasing concentrations of lactate revealed a K_D_ of 293 ± 48 μM and a maximum fluorescence change of 190% (**Figure 2A**). Most single-fluorophore indicators are pH-sensitive, but the pH sensitivity in the physiological range between 7.0 to 7.4 was within experimental error (**Figure 2B**). Next, we performed experiments to evaluate lactate specificity, exposing the purified protein to structurally related monocarboxylates and glucose. None of the tested molecules produced a significant change in the fluorescence response (**Figure 2C**). Next, we evaluated the excitation and emission spectra in the presence or absence of lactate. The results showed both spectra are dependent on lactate, with an isosbestic point at 430 nm and emission peak at 513 nm (**Figure 2D**). This behavior is similar to that of other single-fluorophore indicators developed using the same reporter protein^34^. Since TTHA0766 binds lactate associated with a calcium molecule, a potential interference of calcium cations with the fluorescence lactate response is likely. Thus, we next tested the calcium sensitivity in the hundred micromolar range using purified protein extracts. No calcium sensitivity was detected in the micromolar range. However, complete calcium chelation using EGTA, almost completely abolished the response of the indicator to lactate (**Figure 2E**), confirming the critical role of calcium for lactate binding. This result suggests a possible calcium interference in the nanomolar range, which is consistent with the high affinity of PBP for their ligands and the number of calcium molecules in the lactate binding site of TTHA0766.

**Figure 2.**
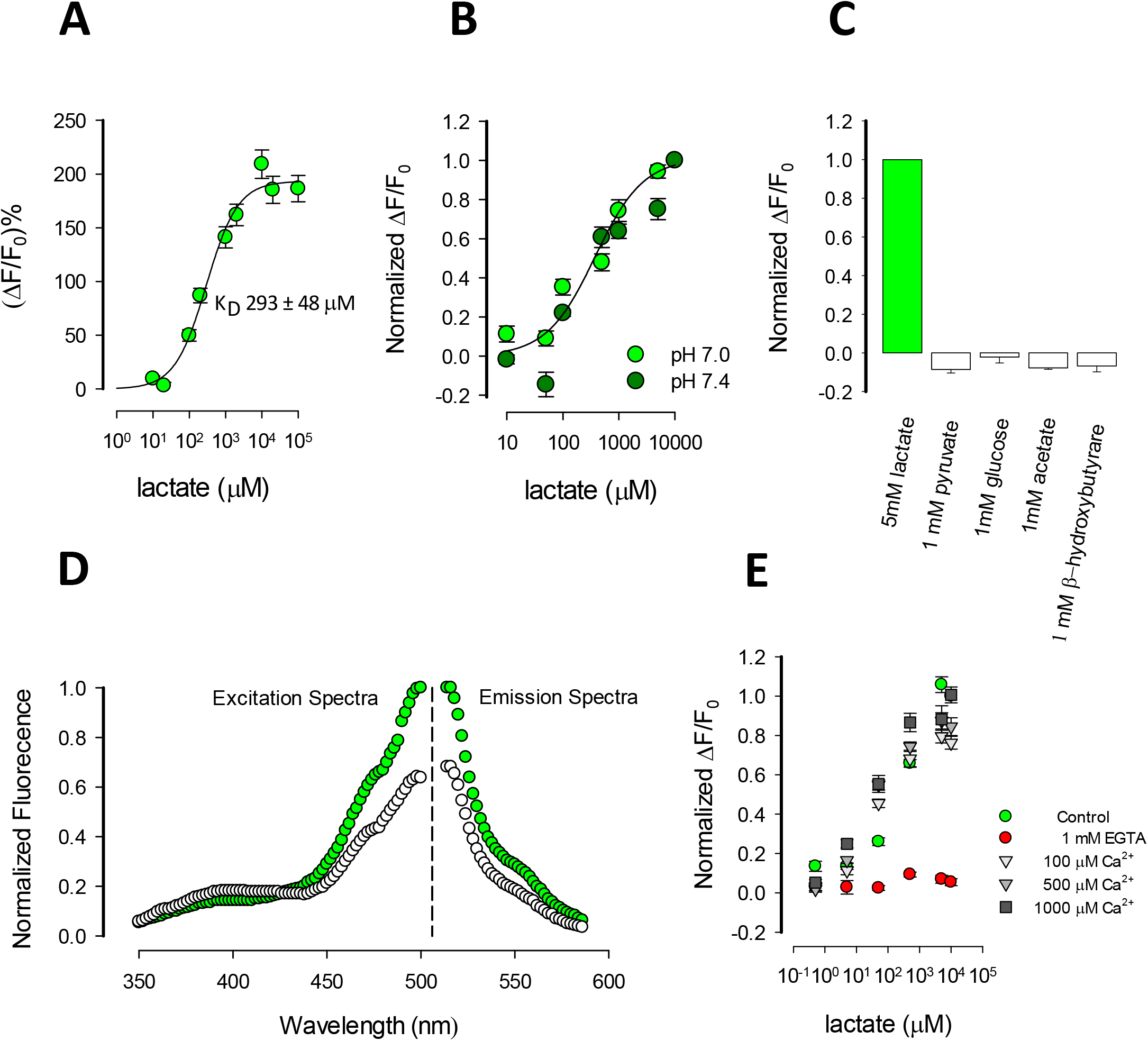
In Vitro Characterization of CanlonicSF. A) Dose-response curve at 10, 20, 100, 200, 1000, 2000, 10000, 20000 and 100000 μM lactate. B) Effect of pH between 7.0 and 7.4 on the dose-response curve of CanlonicSF at 10, 50, 100, 500, 1000, 5000 and 10000 μM lactate. C) CanlonicSF specificity using closely structurally related monocarboxylates and glucose. D) Excitation and emission spectra with (green) and without (white) the presence of 10 mM Lactate. All the experiments were performed using fresh protein extracts from HEK293 cells and are an average of at least 6 replicates wells from two independent protein extracts. E) Calcium interference at 0.5, 5, 50, 500, 5000 and 10000 μM lactate. Nominal calcium concentration in the intracellular buffer was set up using EGTA.

### Characterization of CanlonicSF in mammalian cells

Expressed in mammalian HEK293 cells, the indicator displayed the expected cytosolic distribution with nuclear exclusion (**Figure 3A**). Single-fluorophore indicators are intensiometric, therefore sensitive to volume and focus changes. To minimize such artifacts, we bicistronically expressed mRuby2 (**Figure 3A**), which is insensitive to the analyte. This allowed us to obtain a more robust ratiometric signal. Using HEK293 cells expressing CanlonicSF and mRuby2, a series of experiments were performed to evaluate its *in cellulo* behavior.

**Figure 3.**
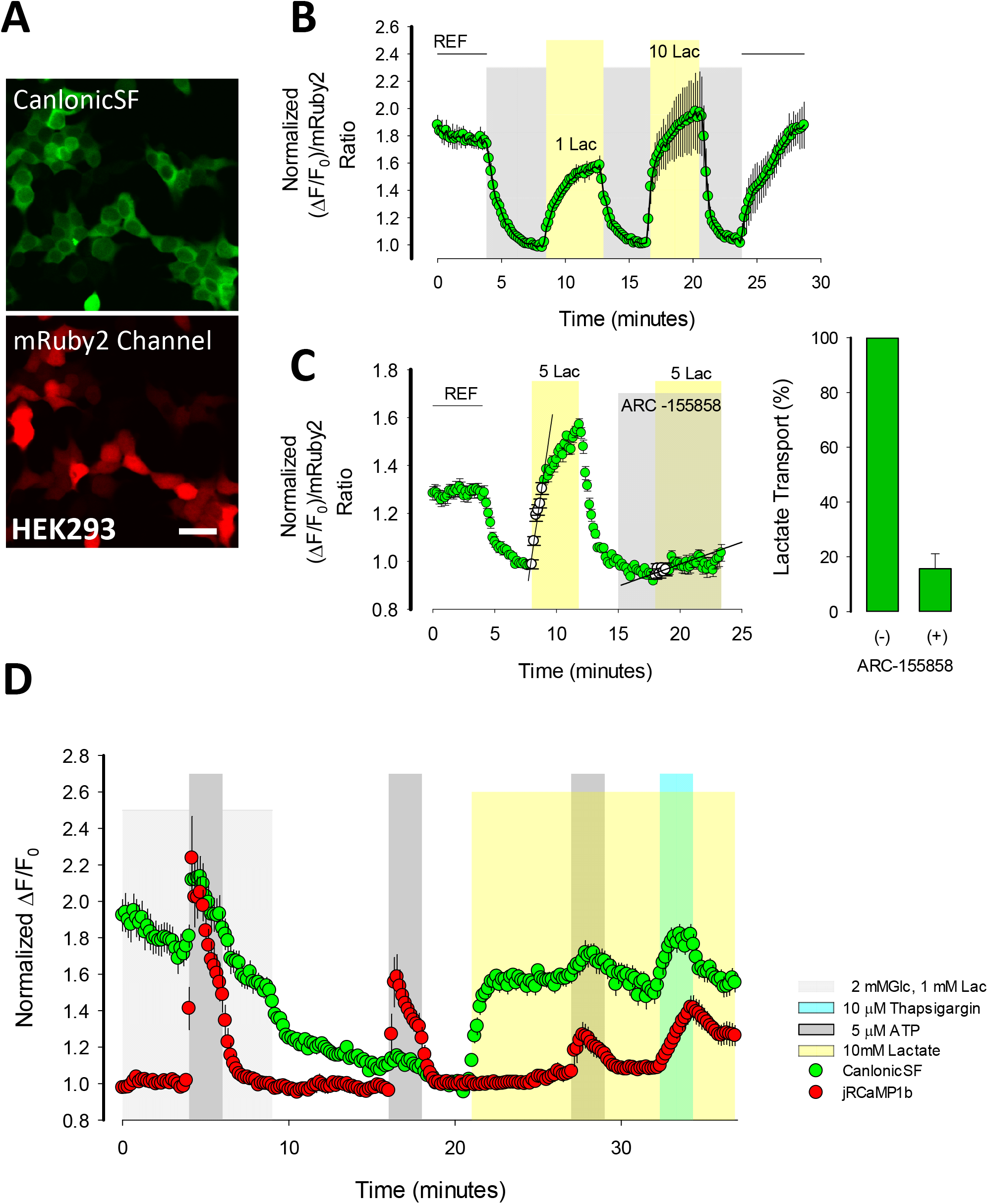
Cytosolic CanlonicSF Behavior in Mammalian Cells. A) Confocal images of HEK293 cells expressing CanlonicSF. Bar represents 50 μm. B) CanlonicSF response expressed in HEK293 cells to 1 and 10 mM lactate pulses. C) MCT transport assay performed in HEK293 cells using 1 μM of ARC-155858 in KRH HEPES buffer. Bar plot is a summary of the percentage of inhibition rates from three independent experiments. D) Cytosolic calcium sensitivity of CanlonicSF expressed in HEK293. Calcium transients were induced using 5 μM ATP and 10 μM thapsigargin. Calcium and lactate multiplexing measurements were performed using CanlonicSF and jRCaMP1b.

Superfused HEK293 cells expressing CanlonicSF were exposed to pulses of increasing lactate concentrations to evaluate the functionality of the indicator. The signal response was fast, consistent with the known permeability of HEK293 cells to monocarboxylates, and it was reversible, permitting before-after experiments (**Figure 3B**). The ΔF/F_0_ was 2.0 which is consistent with our *in vitro* experiments using purified protein extracts.

Single-fluorophore indicators present a higher fluorescent response than FRET-based indicators to a given analyte, making these probes ideal tools to explore transport activity in a cell-based system. To test the suitability of the indicator to evaluate monocarboxylate transporter activity, HEK293 expressing CanlonicSF were exposed to pulses of lactate in the presence and absence of a well-known MCT1 and MCT2 inhibitor ARC-155858^35^. Exposure to 1 μM of ARC-155858 blocked the lactate entry by 84.4% (**Figure 3C**). These results suggest that the CanlonicSF might be an ideal tool to develop a cell-based high-throughput method to screen for MCT inhibitors.

The ligand-binding moiety of CanlonicSF ‒ TTHA0766 ‒ binds calcium-lactate^28^. Experiments using purified protein extracts showed that calcium is needed to detect a lactate-induced fluorescence response and the lactate dose-response curve is not affected by calcium in high micromolar range (**Figure 2E**). Considering the high affinity of PBP for their ligands and the 1:1 stoichiometry of lactate and calcium, we hypothesized that lactate detection is affected by calcium in the nanomolar range. To test this, HEK293 cells expressing CanlonicSF and the calcium red indicator jRCaM1Pb^36^ were treated with 5 μM ATP^37^ and 10 μM of the sarco/endoplasmic reticulum Ca^2+^ ATPase (SERCA) inhibitor thapsigargin^38^ to induced calcium transients. ATP and thapsigargin produced a rapid increase of cytosolic calcium concentration in the nanomolar range, based on the range of response of neuron-expressed jRCaM1Pb to electric stimulation^36^. These calcium transients were accompanied by an increase in the fluorescent signal of the lactate indicator in synchronic fashion (**Figure 3D**). This result confirms that the indicator presents calcium sensitivity and the binding of this cation is necessary for lactate sensing.

To further explore the potential interference of monocarboxylates like acetate, pyruvate, and β-hydroxybutyrate, we designed an *in cellulo* protocol taking advantage of monocarboxylate permeability through the MCTs and trans-acceleration exchange^9^. First, in low lactate concentrations, we pulsed a given monocarboxylate at increasing concentrations and evaluated the induced fluorescent response. We then repeated these pulses in the presence of high lactate and measured the residual lactate as a control signal of monocarboxylate permeability. Exposure to pyruvate, β-hydroxybutyrate, and acetate in low lactate did not increase the fluorescent response (**Figure 4A, C and E**) and in the presence of high lactate, trans-acceleration was detected (**Figure 4B, D and F**). These results support the specificity of CanlonicSF for lactate, based on an approach that maintains an intact intracellular milieu.

**Figure 4.**
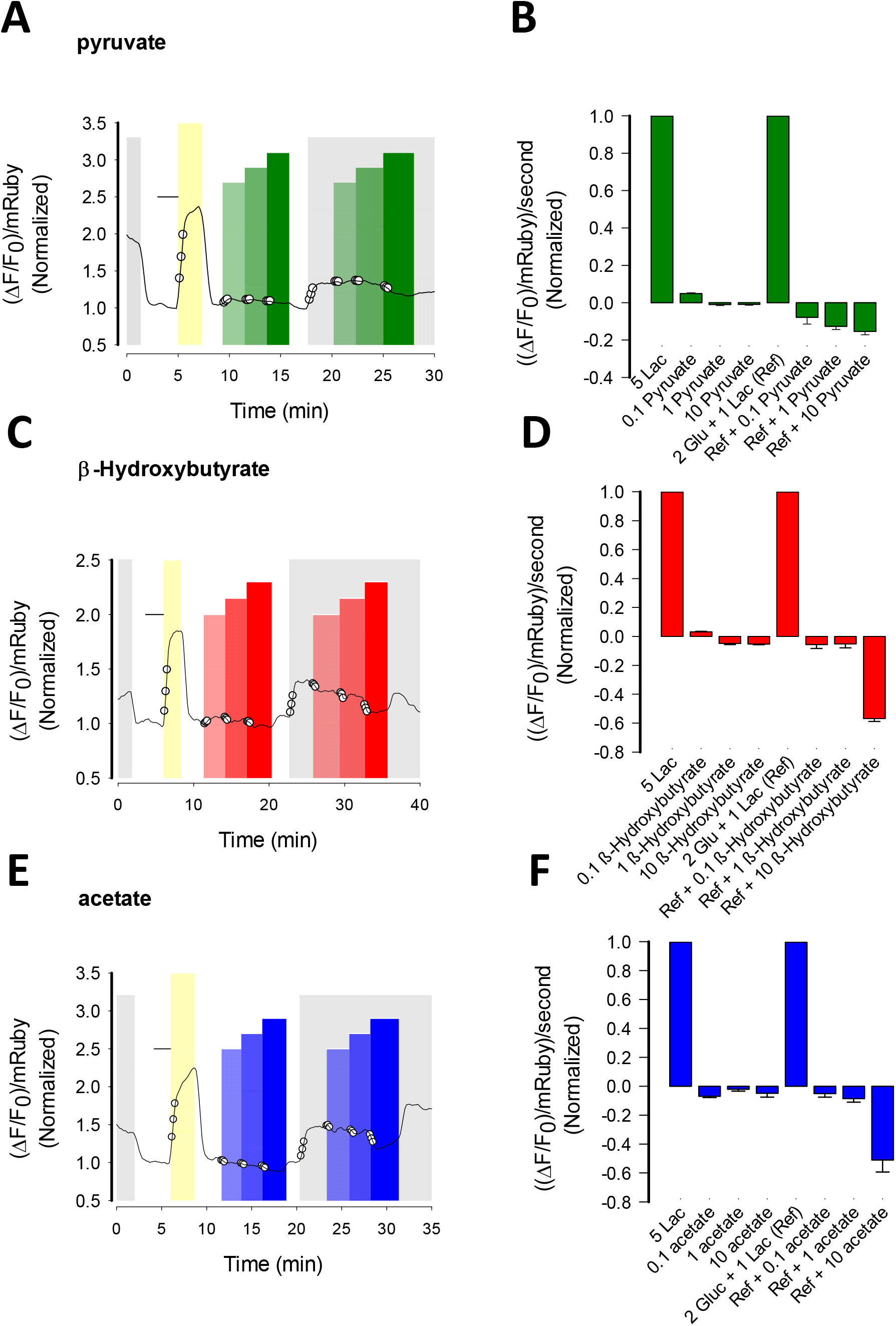
*In cellulo* Specificity Characterization of CanlonicSF. Specificity experiments were performed in HEK293 expressing cytosolic CanlonicSF. A) Pyruvate, C) β-hydroxybutyrate and E) acetate interference protocol consisted in pulses of 0.1, 1, and 10 mM of each substrate in absence of presence of reference KRH bicarbonate buffer with glucose 2 mM and lactate 1 mM. White circles represent the data taken from the experiment to calculate the rate of the fluorescence response. B, D and F) Normalized rate of the fluorescence response induced by pyruvate, β-hydroxybutyrate and acetate, respectively. Data of each set are mean ± s.e.m. (27 cells) from three independent experiments.

### Endoplasmic Reticulum Lactate Dynamics

Based on the *in vitro* experiments using purified protein extracts, CanlonicSF did not present calcium sensitivity in the high micromolar and low millimolar range. This feature points out the potential suitability of the indicator to explore high calcium environments like the endoplasmic reticulum (ER), where the reported calcium concentrations are in that range^39^. We expressed the indicator in COS7 cells with an ER target sequence at the amino-terminus and an ER retention signal at the carboxy-terminus. CanlonicSF displayed a typical reticular pattern (**Figure 5A**) and a response as expected for lactate withdrawal and lactate depletion induced by trans-acceleration exchange using oxamate. The maximal ΔF/F_0_ obtained from F_MIN_ using oxamate, and F_MAX_ reached when blocking the lactate exit to induced CanlonicSF saturation with ARC-155858 was 3.0 (**Figure 5B**).

**Figure 5.**
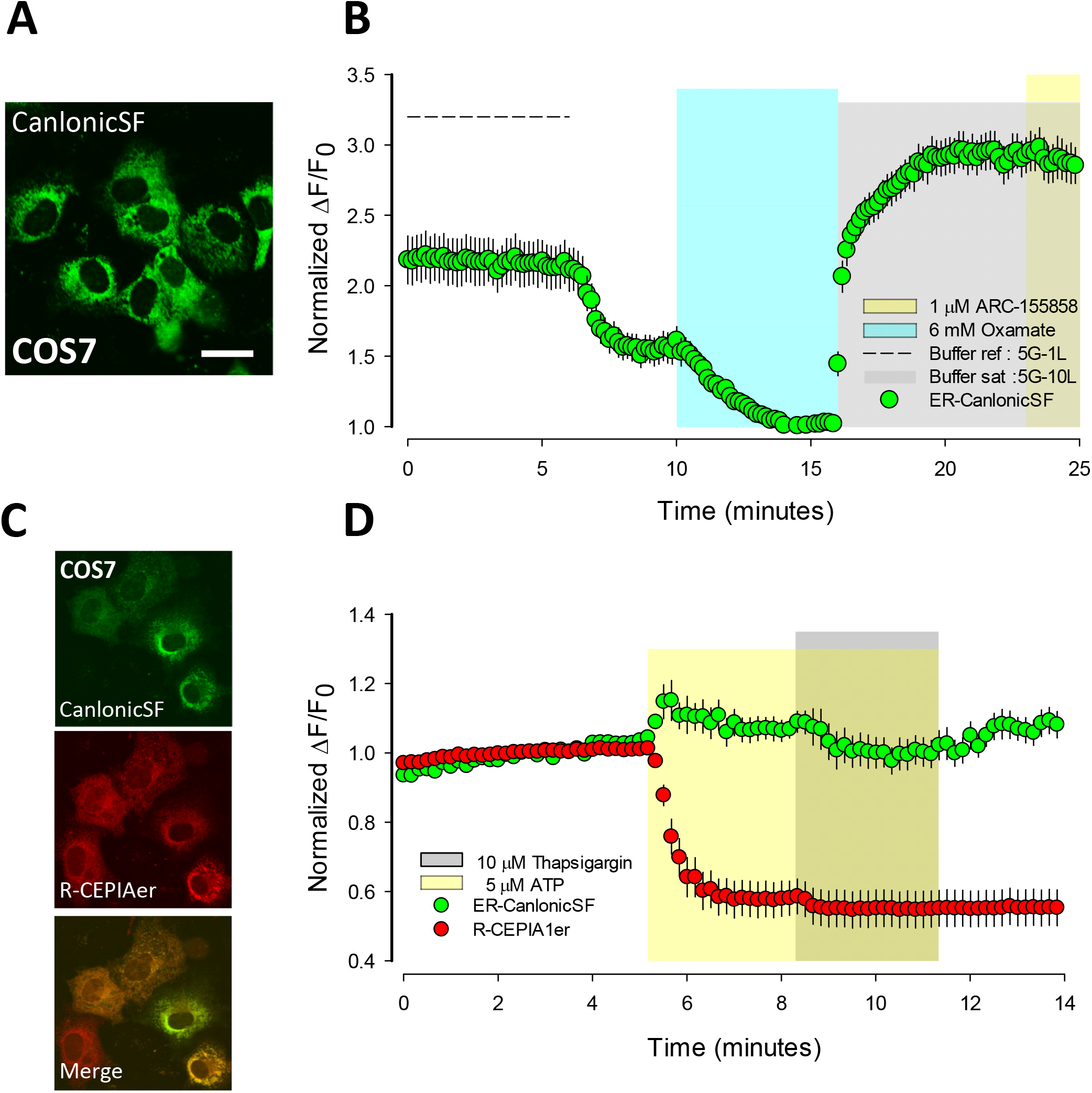
Endoplasmic Reticulum Lactate Dynamics Monitored by CanlonicSF. A) Confocal image of COS7 cells expressing CanlonicSF with ER targeting signal. Bar represents 50 μm. B) Lactate dynamics in ER of COS7 cells. C) Confocal images of COS7 cells expressing CanlonicSF and R-CEPIA1er in the ER. D) Calcium sensitivity of CanlonicSF in the ER using multiplexing measurements of calcium and lactate. Calcium transients were induced using 5 μM ATP and 10 μM thapsigargin. Data are mean ± s.e.m. (15 cells) from one representative experiment.

To test the suitability of the indicator to monitor lactate dynamics in high calcium environments, we performed multiplexing measurements of calcium and lactate using ER targeted red calcium indicator R-CEPIAer^39^ and CanlonicSF, respectively. To do this we co-expressed both indicators in COS7 cells with a target sequence for ER localization. Both indicators displayed a fluorescent signal with a typical reticular pattern. (**Figure 5C**). To induce calcium depletion and therefore calcium mobilization throughout the micromolar range in the ER, we exposed the cells to 5 μM ATP and 10 μM thapsigargin, a well-known pharmacological perturbation that induces calcium depletion in the ER^39^ from the hundred to the dozen on the micromolar range. Calcium depletion reported by R-CEPIAer, monitored using an indicator with K_D_ 368 μM, did not produce any apparent interference in the fluorescent signal of CanlonicSF (**Figure 5D**). Considering the dynamic range of R-CEPIAer, which spans approximately from 3000 to 30 μM, this induced a calcium depletion spanning the micromolar range. Therefore, micromolar calcium did not produce any interference in the indicator signal. The experimental conditions ‒ in terms of carbon sources ‒ did not promote CanlonicSF saturation, therefore we discarded indicator saturation as a plausible explanation for this result. All these experiments support the notion that the single-fluorophore lactate indicator will be instrumental to explore lactate dynamics in high calcium environments.

### Lactate dynamics in *ex vivo* preparations of *Drosophila melanogaster* larval brains

The ability of CanlonicSF to monitor lactate in living tissue was investigated in *Drosophila melanogaster* larvae. To ease buffer access, we studied perineurial glial cells, which present the outermost cell layer of the blood-brain barrier, separating the brain from the surrounding hemolymph. (**Figure 6A**).

**Figure 6.**
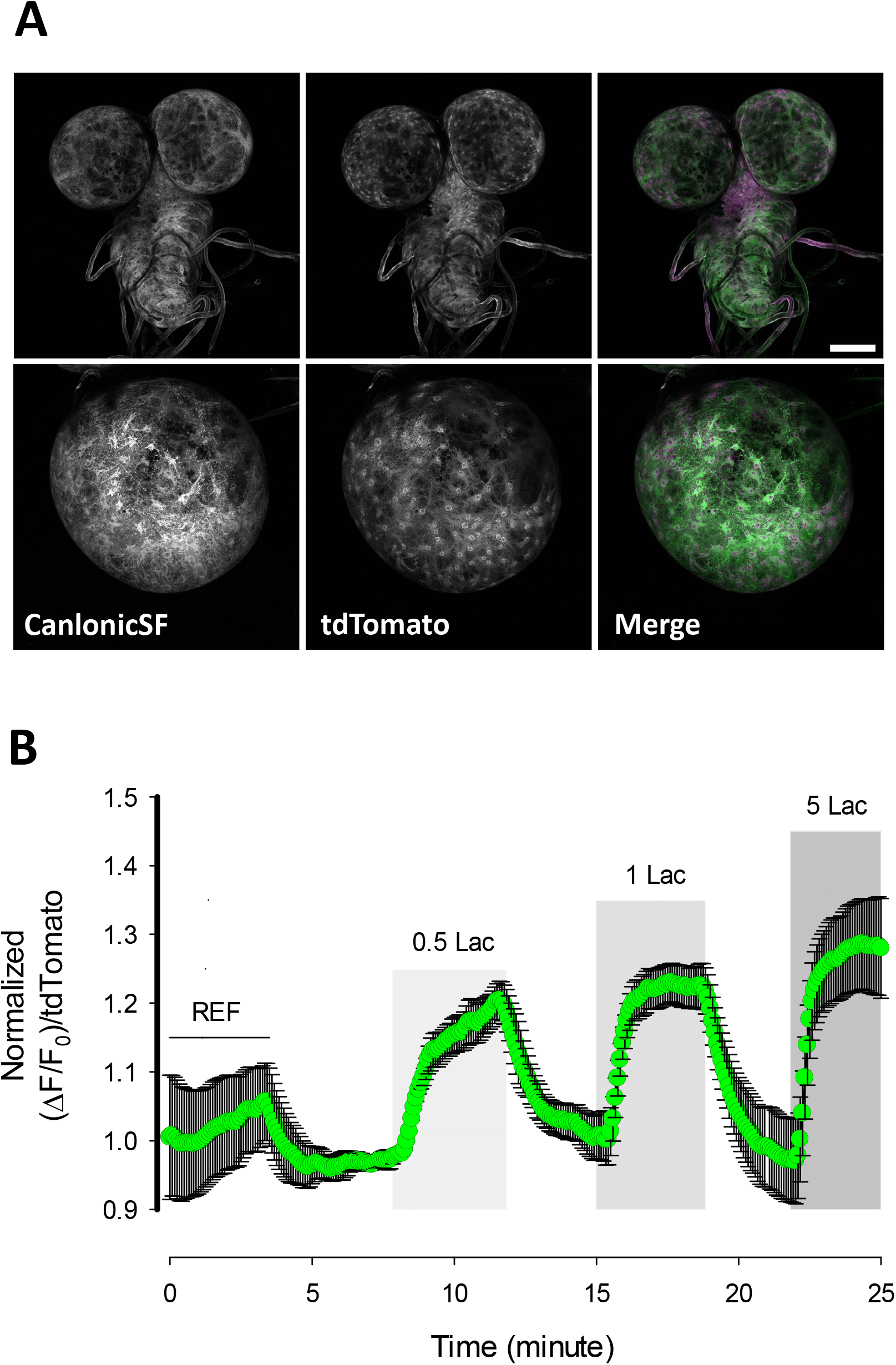
Lactate Dynamics in glial cells of *Drosophila melanogaster* expressing CanlonicSF. Brains were acutely dissected from third instar *Drosophila melanogaster* larvae expressing CanlonicSF in the cytosol of perineurial glial cells. Larval brains were superfused with HL3 buffer containing 5 mM glucose, 1 mM lactate and 0.5 mM pyruvate. A) Direct fluorescence of CanlonicSF and tdTomato in the cytosol of perineurial glial cells (whole CNS (upper panels) and zoom on one optic lobe (lower panels). Scale bar represents 100 μm. (B). Lactate dose-response curve. Shown are mean ± s.e.m. (9 cells) of one representative experiment.

Superfusion of acutely isolated brains with lactate resulted in a quick and reversible increase in cytosolic lactate, revealing the presence of abundant surface lactate transporters in these cells (**Figure 6B**). This result is consistent with the previous results achieved with a single-fluorophore indicator for pyruvate^34^. These experiments show the suitability of the lactate indicator to explore lactate dynamics in *ex vivo* preparations from different model systems.

## DISCUSSION

The present article introduces CanlonicSF as the first genetically encoded single-fluorophore indicator for lactate. This indicator ‒ based on the bacterial PBP TTHA0766 – permits the monitoring of lactate dynamics in mammalian systems and proved to be ideal for subcellular compartments such as ER.

The *in vitro* characterization shows a K_D_ value of 293 ± 48 μM and a dynamic range from 30 to 3000 μM, which span the physiological intracellular lactate concentrations. In spite that we observed variable results in the maximal fluorescence response *in vitro*, we are confident with the reported K_D_, since all the protein extracts showed similar values, independent of their maximal fluorescence response. One-site behavior kinetics allows precise intracellular lactate quantification, which might be more challenging using the current FRET lactate indicator, Laconic. This will be especially critical *in vivo*, where the experimental control is limited. Also, the use of indicators with different K_D_ will allow us to perform a cross-calibration protocol reaching high accuracy in the lactate concentration calculations. Expressed into the cytosol of mammalian cells CanlonicSF showed a homogenous pattern with nuclear exclusion. Transfected cells showed some inclusion bodies due to high expression, but comparable stable cell lines with more controlled protein expression did not show detectable inclusion bodies. The functional experiments showed a reversible fluorescent response to consecutive pulses of increasing lactate concentrations, facilitating our experimental design. The ΔF/F_0_ observed in HEK293 cells is consistent with the measured values obtained using purified extracts. Regarding the specificity, the indicator was exposed to structurally related molecules showing high specificity for lactate, which was confirmed by *in vitro* and *in cellulo* protocols. The main advantage of the *in cellulo* over the *in vitro* approach is the preservation of the intact cellular system, allowing us to discard indirect sources of interference and permits the test protein extract in a more representative environment. We also performed a transport assay that proved that this highly responsive indicator displays better responses and is, therefore, more suitable to set up transport assays in multiwell plate readers for high-throughput screening.

The reported crystallographic structure of TTHA0766 shows a calcium cation in close contact with lactate. Experiments performed to analyze calcium-dependent lactate sensing demonstrated that the presence of calcium is necessary for lactate sensing, although calcium concentrations over 100 μM did not affect the lactate dose-response curve. These results are consistent with the current knowledge about calcium-lactate binding to TTHA0766 and the nanomolar affinity of PBP for their ligands. We conclude that the calcium pocket binding site is fully saturated in the micromolar range.

TTHA0766 binds calcium-lactate. Our *in vitro* experiments showed that the presence of calcium is necessary for lactate sensing and it did not produce any interference with lactate sensing at >100 μM concentrations. These results are consistent with the interference produced by cytosolic calcium transients elicited by 5 μM of ATP and the SERCA inhibitor thapsigargin at 10 μM. Reported cytosolic lactate concentrations are in the low nanomolar range, which together with the known high affinity of PBP for a given ligand explains the impact of cytosolic lactate concentrations on the fluorescent response of the indicator. Furthermore, nanomolar calcium transients did not produce a significant fluorescent response in low lactate, supporting the notion that both calcium and lactate are needed for a proper fluorescent response of CanlonicSF.

Micromolar calcium did not interfere with lactate sensing based on *in vitro* experiments using protein extracts. Therefore, we decided to test the indicator in a high calcium environment like that found in ER, which is reported to be about 400 μM in mammalian cells^39^. The indicator response was smooth and proportional to the observed dynamic range in the cytosol. Pumping out all the residual lactate through trans-acceleration exchange using oxamate and inducing lactate accumulation blocking its exit with ARC-155858, we obtained a robust *in cellulo* maximal ΔF/F_0_ of 3.0. The co-targeting of a red calcium reporter optimized for ER R-CEPIAer and CanlonicSF allowed us to explore potential interference of micromolar calcium on the fluorescent response. Our results are consistent with *in vitro* calcium sensitivity since ER calcium depletion induced by ATP and thapsigargin did not affect the fluorescent response of the lactate indicator. Considering the reported K_D_ for R-CEPIAe, the previous protocol allows to explore calcium interference in the micromolar range. CanlonicSF showed that ER is highly permeable to lactate and the sub-cellular levels in such organelle follows the dynamic has been observed in cytosol. To the best of our knowledge, this is the first time that the lactate dynamics is monitored in ER at single-cell level in an intact system. Also, is the first functional evidence of a lactate transporter in that organelle. All together, these results support the suitability of CanlonicSF to explore lactate dynamics in high calcium environments.

CanlonicSF has room for future improvement. Mutants with lower affinity can be produced by mutating extra amino acid residues in close contact with lactate or the calcium cation. This will be useful for exploring lactate dynamics in tissue or cells that have high lactate concentrations like the muscle. Also, mutations to the linkers might render a new version with a higher maximal fluorescent response; this will be instrumental to develop cell-based methodologies for high-throughput screening of MCT inhibitors. The development of an extracellular version may be generated to explore extracellular lactate dynamics with high spatial resolution, which is not currently possible with available techniques.

In summary, we developed a calcium-lactate single fluorophore indicator based on the PBP bacterial protein TTHA0766. We envision that this calcium-dependent lactate indicator will be of practical interest in the exploration of lactate dynamics in high calcium environments like the ER in both *in vitro* and *ex vivo* preparations, and the development of high-throughput methodologies for screening of MCT inhibitors.

## ACKNOWLEDGEMENTS

This work was supported by FONDECYT grant Initiation into Research number 11150930 and joint grant from the Agencia Nacional de Investigacion y Desarrollo (ANID)-Chile and the Bundesministerium für Bildung und Forschung (BMBF)-Germany (ANID-BMBF 180045) and The Centro de Estudios Científicos (CECs) is funded by the Chilean Government through the Centers of Excellence Basal Financing Program of ANID.

